# Fourfold increase in photocurrent generation of *Synechocystis* sp. PCC 6803 by exopolysaccharide deprivation

**DOI:** 10.1101/2024.02.09.579478

**Authors:** Laura T. Wey, Evan Wroe, Viktor Sadilek, Linying Shang, Xiaolong Chen, Jenny Z. Zhang, Christopher J. Howe

## Abstract

Photosynthetic microorganisms, including algae and cyanobacteria, export electrons in a light-stimulated phenomenon called ‘exoelectrogenesis’. However, the route(s) by which electrons reach an external electrode from the cell remain(s) unclear. For the model cyanobacterium *Synechocystis* sp. PCC 6803, it has been established that electron transfer does not depend on direct extracellular electron transfer by type IV pili. However, the role of the exopolysaccharide matrix in which cells are embedded has not been investigated. We show that a *Synechocystis* mutant with substantially reduced exopolysaccharide production has a four-fold greater photocurrent than wild-type cells. This increase is due in part to increased adhesion of exopolysaccharide-deficient cells to electrodes. Stirred system experiments reveal that a substantial portion of the photocurrent depends on an endogenous diffusible electron mediator, supporting indirect extracellular electron transfer as the bioelectrochemical mechanism of exoelectrogenesis. These findings will be important in harnessing exoelectrogenesis for sustainable electricity generation in biophotovoltaic devices.

## Introduction

Photosynthetic microorganisms, including algae and cyanobacteria such as *Synechocystis* sp. PCC 6803 (hereafter referred to as *Synechocystis*), can export electrons in a light-stimulated phenomenon called ‘exoelectrogenesis’, which has enormous potential applications for sustainable generation of electricity from light and water in biophotovoltaic devices (BPVs).^1^ However, the bioelectrochemical mechanism of electron export is poorly understood, hindering efforts to increase BPV outputs.

Based on studies of electron export by heterotrophic electrogenic microorganisms, two broad classes of physiological mechanisms of extracellular electron transfer (EET) from microorganisms to electrodes can be proposed.^2^ Direct extracellular electron transfer (DEET) involves direct contact and electron transfer between the cell and the electrode via conductive ‘microbial nanowire’ extracellular appendages, redox proteins on the cell surface, or redox proteins or metabolites embedded in the extracellular matrix. Indirect extracellular electron transfer (IEET) involves electron mediators diffusing from the cell to the electrode. The mediators could be continuously produced metabolites that are oxidised at the electrode surface, or metabolites that cycle between the cell and the electrode. Exogenous electron mediators can also be added.

The nanowires of heterotrophic bacteria such as *Geobacter sp*. or *S. oneidensis MR-1* are believed to be pili or filaments including multi-haem cytochromes.^3–5^ There is some evidence of direct extracellular electron transfer (DEET) in *Synechocystis* via extracellular appendages called type IV pili,^6,7^ but more recent work disagrees.^8,9^ The outer membrane and periplasmic space of *Synechocystis* have been shown to have a role in gating exoelectrogenesis.^10^ There is no biochemical or genomic evidence for the existence in cyanobacteria of the multihaem cytochromes found in nanowires of *Geobacter sp*. or *S. oneidensis MR-1* (and which are also located at the cell surface).

A variety of different endogenous electron mediators facilitate IEET by heterotrophic electrogens. In *Shewanella sp.*, the secretion of flavin mononucleotide and riboflavin contributes greatly to their EET.^11–14^ Flavins also mediate IEET in hundreds of Gram-positive bacterial species within the Firmicutes.^15^ However, there is no genomic evidence for systems for secretion of flavins in cyanobacteria.^2^ Phenazines are another class of endogenous electron mediator produced by bacteria such as pseudomonads, including *Pseudomonas aeruginosa.*^16^ However, they are not naturally produced by cyanobacteria,^17^ (although *Synechocystis* has been genetically engineered to biosynthesise the phenazine pyocyanin).^18^ The final class of endogenous electron mediator studied from electrogens are quinones. *Escherichia coli* uses a hydroquinone derivative as endogenous electron mediator.^19^ *Shewanella oneidensis* MR-1 uses 2-amino-3-carboxy-1,4-naphthoquinone (ACNQ) as an electron shuttle, with the cells releasing a precursor structure 1,4-dihydroxy-2-naphthoic acid (DHNA) into the environment where it spontaneously converts into ACNQ.^20,21^ The phylloquinone biosynthetic pathway of cyanobacteria involves DHNA as an intermediate.^22^ Benzoquinones are common electron mediators found naturally in ETCs, such as plastoquinone in the photosynthetic and respiratory electron transport chains in cyanobacteria.^23^ This raises the possibility that quinones might function in IEET in cyanobacteria.

There is some evidence from cyclic voltammograms (CVs) of cyanobacterial biofilms of an endogenous electron mediator that enables IEET from the cell to an electrode (**Fig. 1a**).^24,25^ In principle, this could simply diffuse or be transported from the cell in a reduced form and transfer electrons to the electrode. Alternatively, it could exist outside the cytoplasm and accept electrons from a reductase in the cytoplasmic or outer membrane. Kusama *et al*. (2022) achieved an enhancement in photocurrent generation of *Synechocystis* by outer membrane deprivation, consistent with the mechanism of a mediator diffusing across the outer membrane (and possibly the inner membrane as well).^10^ Based on redox midpoint potential, Zhang *et al*. (2018) suggested the mediator from *Synechocystis* was a benzoquinone or flavin derivative.^24^ Saper *et al*. (2018) suggested it was a temperature-sensitive and water-soluble quinone, flavonoid or small peptide.^25^ The mediator was recently proposed to be NADPH by Shlosberg *et al.* (2021), who identified NADPH using fluorescence spectroscopy in the electrolyte after photoelectrochemistry experiments, and showed that exogenous addition of NADPH increased photocurrent output from *Synechocystis* cells on graphite electrodes.^26^ Conversely, using indium tin oxide electrodes, Kusama *et al*. (2022) observed different fluorescence spectra between NADPH and the electrolyte, and found that the intracellular NADPH signal did not change when photocurrent was increased.^10^ Therefore, the identity of all the endogenous electron mediators is the subject of debate at present.^27^

**Fig. 1.**
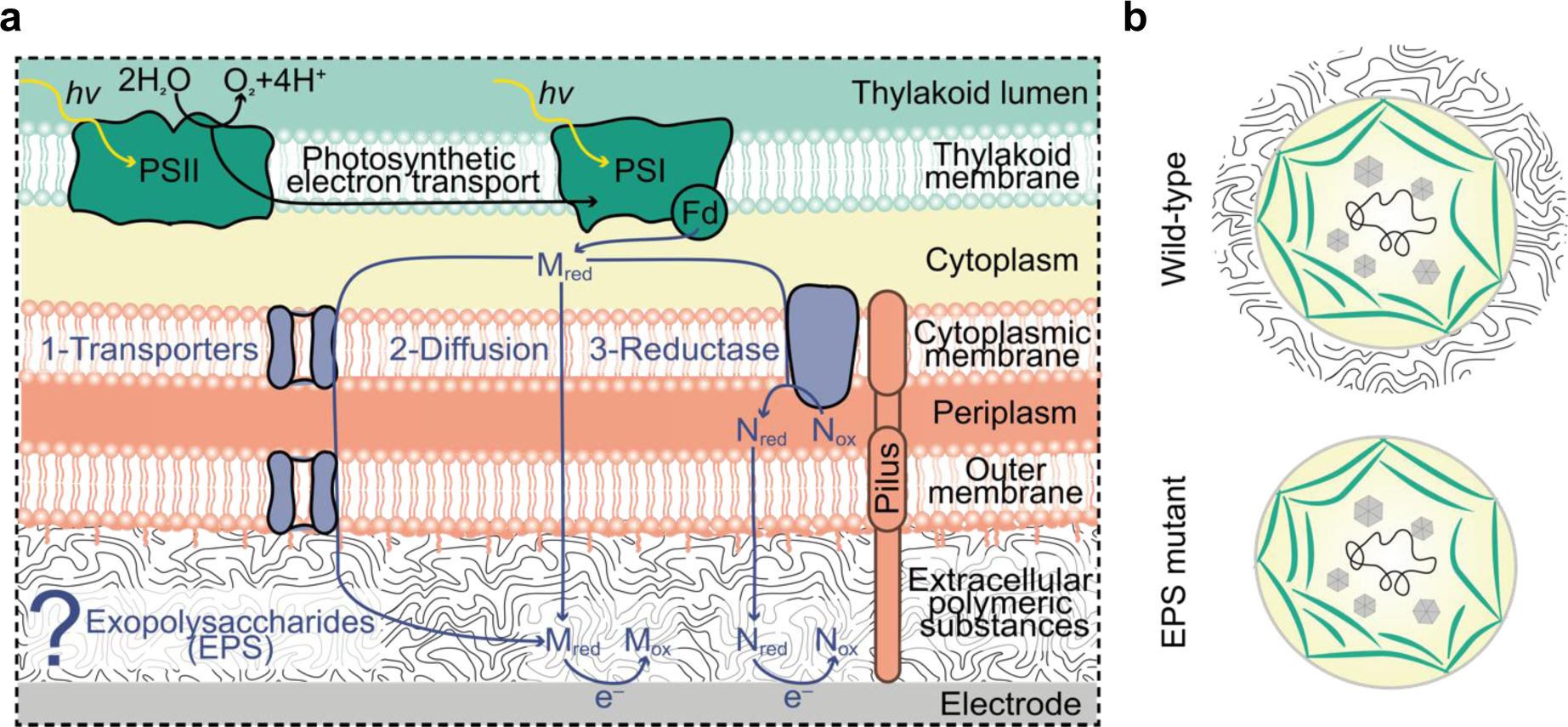
Exoelectrogenesis and exopolysaccharides in *Synechocystis sp.* PCC 6803 cells. **a** Diagram of present understanding of exoelectrogenesis in *Synechocystis sp.* PCC 6803 cells. Shows a simplified photosynthetic electron transport chain in the thylakoid membrane. ‘M’ and ‘N’ are endogenous electron mediators produced and exported by the cells under illumination. The endogenous mediator M is hypothesised to exit the cell via transporters or diffusion, or transfer electrons via reductase(s) coupled with another mediator, N (i.e. one mediator reduced by the photosynthetic electron transport chain diffuses to the cytoplasmic face of the cytoplasmic membrane, where a putative reductase oxidises it and simultaneously reduces another mediator in the periplasm that then exits the cell). PSII – photosystem II, PSI – photosystem I. **b** Diagram of wild-type and exopolysaccharide (EPS) mutant cells.

The secreted extracellular polymeric substances that constitute the matrix in which cells in a biofilm are embedded may also mediate electron transfer. This matrix consists of exopolysaccharides (EPS), proteins, humic substances, nanostructured membrane vesicles and other macromolecules.^28^ Heterotrophic exoelectrogens such as *Geobacter sp.*, *Shewanella oneidensis* and *Pseudomonas putida* are known to have redox metabolites and proteins around their EPS.^29,30^ In photosynthetic exoelectrogens, the role of the EPS in exoelectrogenesis has not been explored yet.

*Synechocystis* EPS comprise 12 different monosaccharides, and are either closely associated with the cell surfaces (capsular polysaccharide) or released into the environment (released polysaccharide).^31,32^ The EPS are decorated with sulphate and uronic acid which confer anionic character on them.^33^ Acetylations, pyruvylations and peptide modifications are also present and modulate hydrophobicity.^33^ The EPS protects cells from high salinity and osmotic stress,^28,34,35^ which also could be involved in exoelectrogenesis.^36–38^ It has been shown that the *Synechocystis* EPS is effective at chelating metal ions to protect cells from heavy-metal toxicity.^28,39,40^ These ions could also play a role in exoelectrogenesis as redox carriers.^41,42,10^

To investigate the role of the EPS in the mechanism (DEET or IEET) of exoelectrogenesis, we analysed the photoelectrochemical performance of a *Synechocystis* double mutant with significantly decreased EPS, which was previously generated and characterised by Jittawuttipoka *et al*. (2013) (**Fig. 1b**).^28^ The EPS mutant has an interrupted *slr1875* gene, which has homology to the *exoD* gene of *Rhizobium meliloti*, where it is suspected to play a role in exopolysaccharide synthesis.^43^ The EPS mutant also has an interrupted *sll1581 gene*, which has homology to *gumB* from *Xanthamonas axonopodis* which encodes a known exopolysaccharide exporter.^44^ The EPS mutant has been previously shown to produce 11-fold lower amounts of capsular polysaccharides attached to cells and 2-fold lower amounts of polysaccharides released in the liquid culture medium than wild-type cells.^28^

The EPS-deficient cells were found to have strikingly increased exoelectrogenic activity. The physical interactions of the EPS mutant cells with the electrode and between cells were also analysed to test if EPS affects cell loading on electrodes. These physical interactions are important as they may also influence the efficiency of the electrical connection between the cells and the electrodes (‘wiring’). Experiments combining photoelectrochemistry experiments with convection provided further evidence of a diffusible electron mediator. Cells with significantly reduced EPS were found to be better interfaced with the electrode, and studies using a fluorescent probe suggested the EPS hinders the ability of molecules such as redox mediators to move in the vicinity of wild-type cells.

## Results

### Mutant with less exopolysaccharide yields higher photocurrent

To investigate whether the EPS affect exoelectrogenesis in *Synechocystis*, the photoelectrochemical performance of the EPS mutant was compared with the wild-type strain from which it was derived. Suspensions of EPS mutant and wild-type cells concentrated to 37.5 nmol of chlorophyll per sample were loaded onto hierarchically structured inverse-opal indium-tin oxide (IO-ITO) electrodes. The high surface area of these electrodes boosts cell loading and electrochemical signals^24,45^ Chronoamperometry was conducted at an applied potential of +0.1 V vs Ag/AgCl (saturated KCl, equivalent to +0.3 V vs Standard Hydrogen Electrode (SHE)). This applied potential yields the maximum photocurrent for wild-type *Synechocystis* cells as shown from stepped chronoamperometry in a previous study also using IO-ITO electrodes.^24^ During chronoamperometry, the loaded electrodes were exposed to cycles of 60 s of 680 nm light at 50 µmol_photons_ m^−2^ s^−1^ (approximately 1 mW cm^-2^ equivalent) to drive photosynthesis, followed by 90 s of dark (after which periods the steady-state currents in the light and dark had stabilised).

Both wild-type and EPS mutant cells showed a complex current output over time under light/dark cycles (the photocurrent profile) with a pattern of features including peaks and troughs before reaching a steady state (**Fig. 2a, Supplementary Fig. 1**). The photocurrent profiles of wild-type and EPS mutant cells showed two peaks at approximately 5 s and 25 s after illumination, a trough that reached its minimum approximately 12 s after illumination, and a broader trough that reached its minimum approximately 15 s after the light was turned off. These photocurrent profiles were similar to previous results using a different wild-type *Synechocystis* strain also loaded on IO-ITO electrodes.^9,24^ It has been previously found that isolated photosystem II and spheroplasts without an intact periplasmic space and outer membrane with EPS do not yield complex photocurrent profiles,^9,24^ suggesting that the complexity in the photocurrent profile of cyanobacterial cells lies in the mechanism of exoelectrogenesis across the outer layers of the cell structure. The presence of similar features in the photocurrent profiles of wild-type and EPS mutant cells therefore suggests that the EPS is not itself an essential component of exoelectrogenesis.

**Fig. 2.**
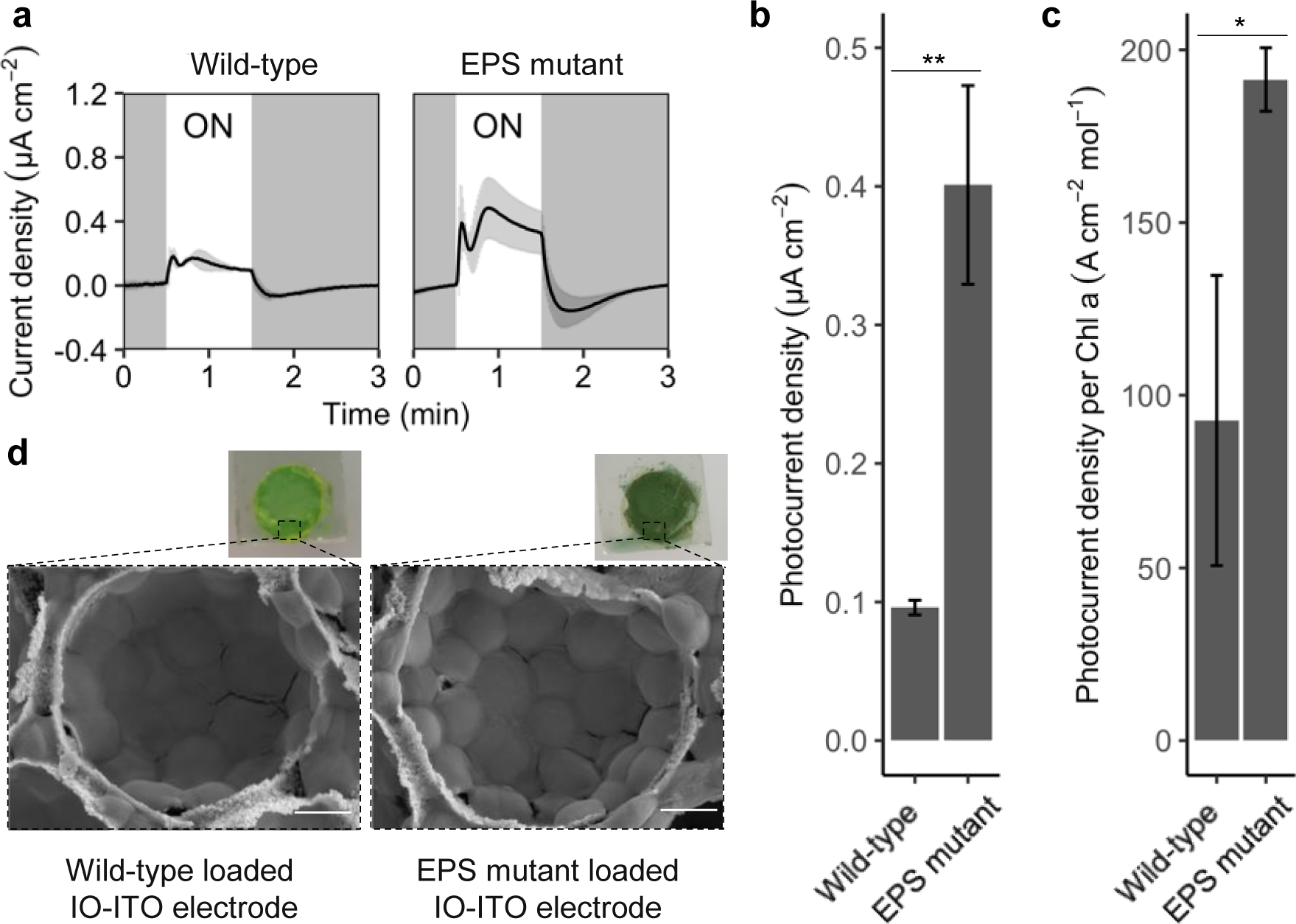
Photoelectrochemical performance of exopolysaccharide mutant and background wild-type cells. **a** Photocurrent profiles where ON = light on and grey panels = light off. The current was normalised to the geometric area of the IO-ITO electrode to obtain current densities. The same data with different y-axis scales are in Supplementary Fig. 1. **b** Photocurrent outputs calculated from photocurrent profiles in panel A. **c** Photocurrent outputs normalised to chlorophyll loading. Data presented as mean ± standard error of the mean of 3-4 biological replicates, ** = P < 0.01 and * = P < 0.05 using an unpaired t-test. **d** Photographs of loaded IO-ITO electrodes after photoelectrochemistry experiments, and scanning electron microscope (SEM) images of IO-ITO electrodes loaded as per the protocol for photoelectrochemistry experiments and then rinsed in BG11 medium. Geometric surface area of the electrode in the photographs is 0.79 cm^2^. Scale bar is 2 µm in SEM images.

The magnitudes of the photocurrent profiles of wild-type and EPS mutant cells were different although the electrodes had been loaded with cells containing the same amount of photosynthetic material. The photocurrent output was calculated as the difference between the steady-state currents in the light and dark (**Supplementary Fig. 2**). The EPS-less mutant yielded four-fold higher photocurrent output (0.40 ± 0.07 µA cm^-2^, n = 4) than wild-type cells (0.096 ± 0.005 µA cm^-2^, n = 3, P = 0.004) (**Fig. 2b**).

Although suspensions of wild-type and EPS mutant cells containing the same total amount of chlorophyll had been loaded onto IO-ITO electrodes before the photoelectrochemistry experiments, the electrodes appeared more densely loaded with EPS mutant cells than with wild-type cells after the experiments (**Fig. 2d**), although both the EPS mutant and wild-type cells appeared intact on the IO-ITO electrodes as visualised by scanning electron microscopy (**Fig. 2d**).

Therefore, the amount of chlorophyll remaining on the electrodes after photoelectrochemistry experiments was measured to normalise photocurrent outputs to the amount of photosynthetic material (for which chlorophyll is a proxy). Surprisingly, the chlorophyll amount remaining on electrodes loaded with EPS mutant cells was approximately double that on electrodes loaded with wild-type cells (wild-type: 1.7 ± 0.7 nmol^-1^_Chl_, EPS mutant: 4.4 ± 1.0 mol^-1^_Chl_). When the photocurrent outputs were normalised to the chlorophyll loading, the difference in output between EPS mutant and wild-type was decreased, but the EPS mutant still yielded two-fold higher photocurrent output (191 ± 9 A cm^-2^ mol^-1^_Chl_, n = 4) than wild-type cells (93 ± 42 A cm^-2^ mol^-1^_Chl_, n = 3, P = 0.04) (**Fig. 2c**). Given that cells containing equal amounts of chlorophyll had been loaded at the start, the difference at the end of the experiment indicates that the EPS mutant cells were better able than the wild-type cells to remain inside the 3D-structure of the IO-ITO electrodes when submerged in the BG11 electrolyte.

### Mutant with less exopolysaccharides has greater cell-electrode and cell-cell interactions

It was hypothesised that the EPS mutant had altered physical interactions with the electrode and other cells compared to the wild-type cells, that affected loading onto the electrodes (**Fig. 2d** photographs) and contributed to some of the difference in the photocurrent outputs prior to normalisation by chlorophyll amount. Therefore, we assessed the cell-electrode and cell-cell interactions in a number of ways.

The formation of biofilms on ITO by EPS mutant and wild-type cells was assessed using ITO coated PET (ITO-PET) with a flat structure in a crystal violet assay. The EPS mutant cells appeared to form more uniform biofilms than wild-type cells (**Fig. 3a**), the EPS mutant cells had significantly greater biofilm formation on unmodified ITO-PET than wild-type cells (WT: Abs_600_ = 0.44 ± 0.09, EPS-less: Abs_600_ = 0.59 ± 0.01, n = 4-6, P = 0.009) **Fig. 3b**). The biofilm formation assay was repeated with ITO-PET electrodes roughened with fine sandpaper. Increasing the roughness of the ITO-PET with more treatment with sandpaper had no effect on the density of biofilms formed by wild-type cells. However, the EPS-less cells had significantly greater biofilm formation on rougher ITO-PET than ITO-PET that had not been roughened.

**Fig. 3.**
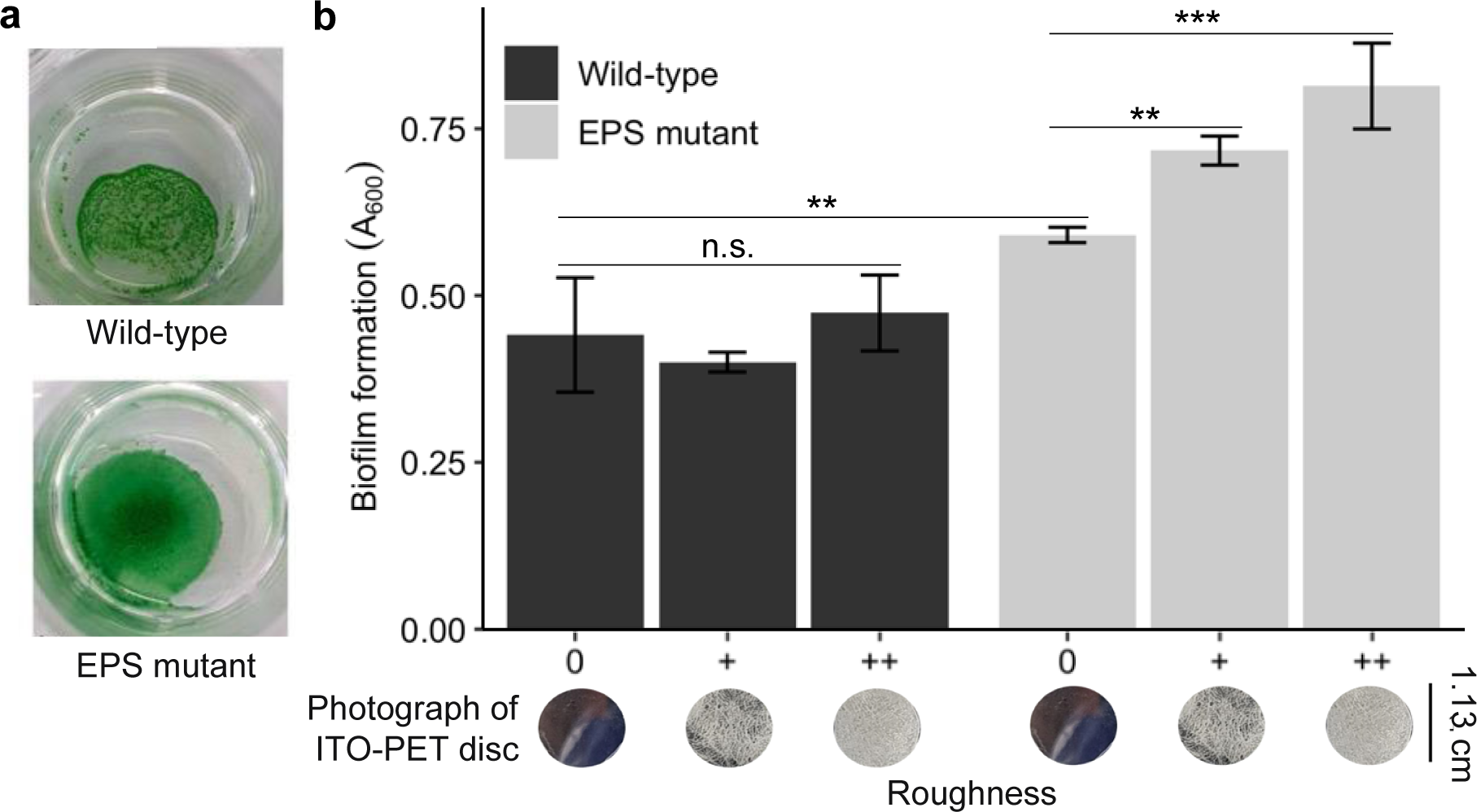
Biofilm formation of exopolysaccharide mutant and background wild-type cells. **a** Representative photographs of biofilms on flat ITO-PET after the final washing step had been carried out during a biofilm formation assay, showing the more uniform appearance of the biofilms with mutant cells. **b** Crystal violet assay of biofilm formation of cells loaded as per the protocol for photoelectrochemistry on ITO-PET electrodes unmodified and roughened. Roughness: 0 (unmodified ITO-PET), + (ITO-PET with low roughness from some sandpaper treatment), ++ (ITO-PET with high roughness from more sandpaper treatment). Data presented as mean ± standard deviation of 4-6 replicates, n.s. = not significant, ** = P < 0.01, *** = P < 0.001 using a one-way ANOVA and post hoc Tukey HSD test.

The cell-cell interaction of EPS mutant and wild-type cells was measured using an assay of their flocculation (i.e. aggregation of cells in suspension). Cells in early stationary phase (as per the protocol for photoelectrochemistry and biofilm assays) and in exponential phase were treated with gentle shaking and under constant illumination with moderate light (40 µmol_photons_ m^-2^ s^-1^) for 48 h, then their flocculation was observed and quantified by measuring the intensity of pixels in photographs, with pixels with increased intensity indicating increased flocculation.^46^ The wild-type cells in exponential phase appeared not to flocculate (**Fig. 4a**). EPS mutant cells in exponential phase had significantly greater flocculation (0.050 ± 0.002 a.u., n = 3, P = 0.03) than wild-type cells in exponential phase (0.038 ± 0.005 a.u.) (**Fig. 4b**). No difference was seen in the flocculation of cells in early stationary phase. This was consistent with previous findings that the EPS mutant cells sedimented, whereas the wild-type cells remained dispersed in liquid medium.^28^

**Fig. 4.**
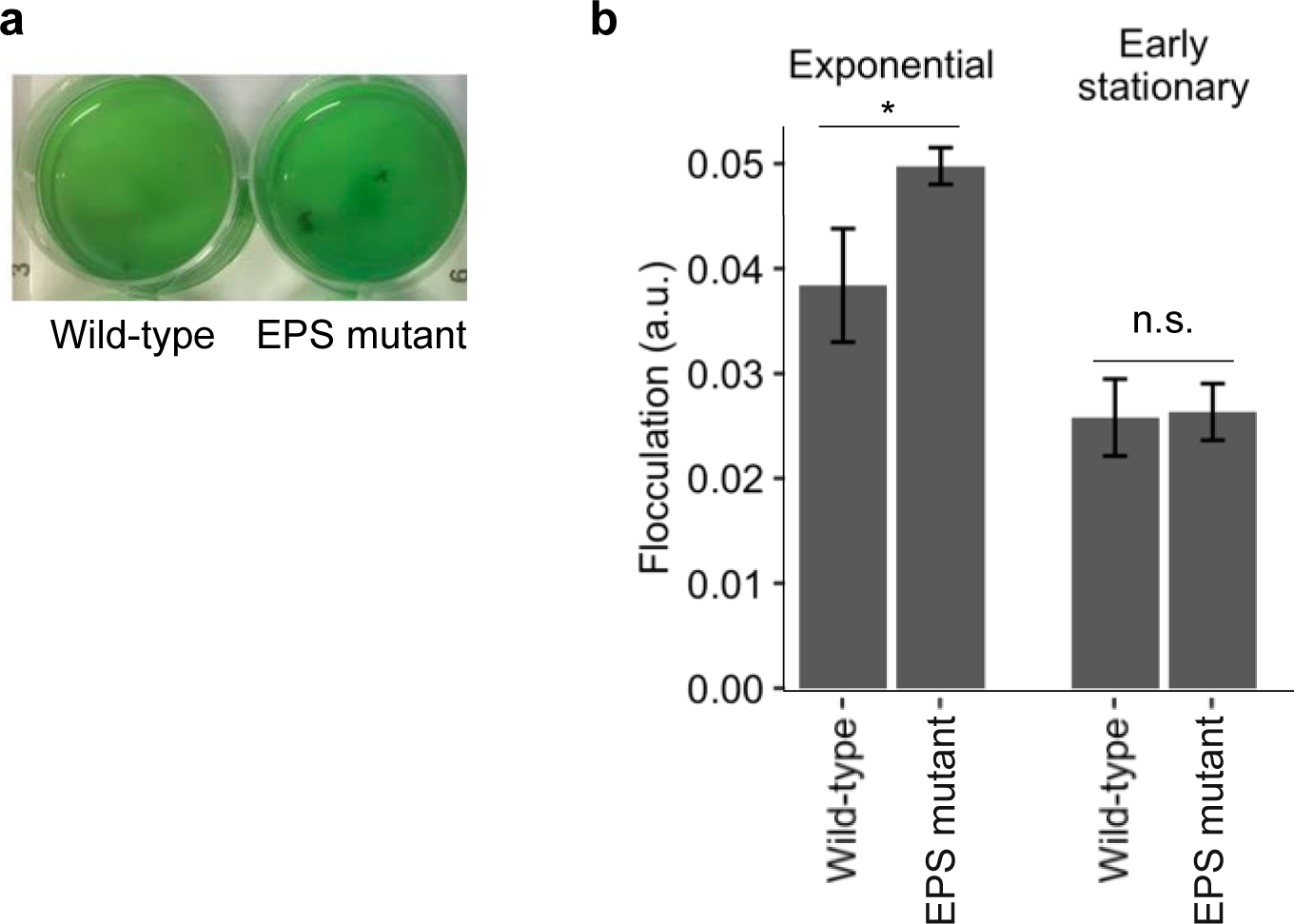
Flocculation of exopolysaccharide mutant and background wild-type cells. **a** Photographs of exponential cultures in multi-well plates after a flocculation assay. **b** Quantification of flocculation. Data presented as mean ± standard deviation of three biological replicates, n.s. = not significant, * = P < 0.05 using a one-way ANOVA and post hoc Tukey HSD test.

The packing of the cells in biofilms on electrodes was visualised using light microscopy. Equal volumes of wild-type or EPS mutant cells from early stationary phase cultures (200 μl, OD_750_ = 1.2) were dropcast onto an ITO-coated glass coverslip. Cells were left for 18 hours in the dark to settle and form a biofilm, which was similar to the protocol for photoelectrochemistry experiments. The biofilms were visualised using light microscopy (**Fig. 5a**). The EPS mutant cells were observed to pack together more closely than wild-type cells (cell-to-cell distance wild-type: 1.33 ± 0.3 µm, EPS mutant: 0.38 ± 0.07 µm, P = 0.007, n = 3 i.e. three cultures each measuring 10 cells) than wild-type cells (**Fig. 5b**). The EPS mutant cells were also observed to be in the same focal plane of the light microscope (i.e. had settled at the same height on the coverslip), whereas the wild-type cells were observed to be distributed over a wider vertical range of the biofilm.

**Fig. 5.**
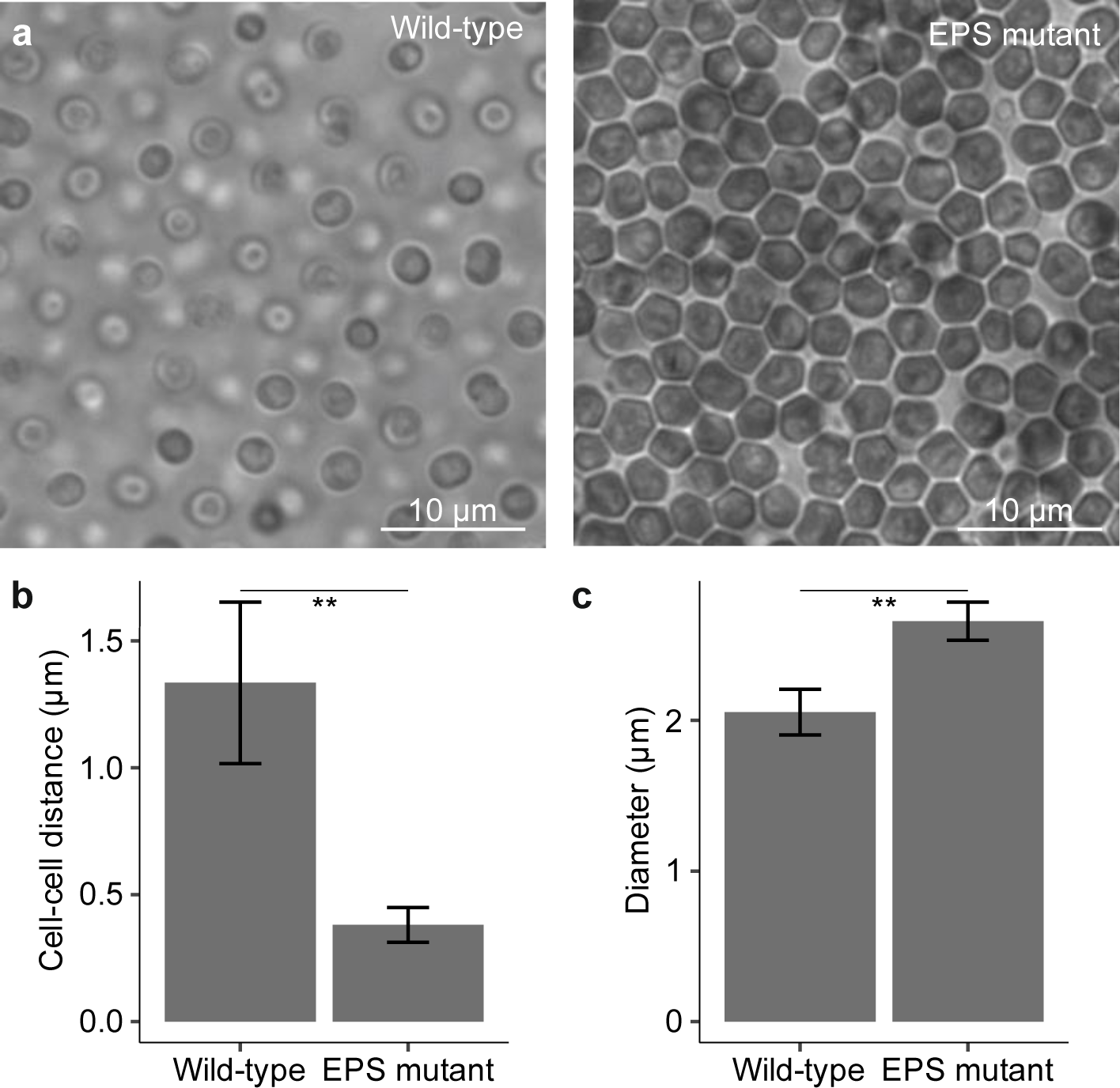
Dimensions of exopolysaccharide mutant and background wild-type cells packing in monolayers. **a** Bright field microscopy images of biofilm of cells. Images representative of 3 biological replicates. Scale bar = 10 µm. **b** Cell to cell distance. **c** Cell diameter. Data presented as the mean ± standard deviation of 30 cells (3 biological replicates each measuring 10 cells, and errors propagated, n = 3 used for statistical analysis), *** = P < 0.001 using a student’s t-test.

The EPS mutant cells had an approximately 1.3-fold greater diameter than wild-type cells (wild-type: 2.06 ± 0.15 μm, EPS mutant: 2.66 ± 0.13 μm, P = 0.0006, n = 3 i.e. three cultures each measuring 10 cells) (**Fig. 5c**). To investigate whether this difference in cell size could explain the disparity in photocurrent generation between wild-type and EPS mutant cells, we conducted photoelectrochemical tests on two different-sized strains of ‘wild-type’ *Synechocystis*. These strains, despite varying in cell sizes by a similar ratio to that measured between wild-type and EPS mutant cells, exhibited similar photocurrent output (**Supplementary Fig. 4**). This suggests that the differences in photocurrent generation between wild-type and EPS mutant cells are not due to the differences in their size.

These results indicate that the EPS mutant cells interacted more closely with the electrode and with each other, forming a stronger biofilm. An increased amount of biofilm would account for some of the increased photocurrent output of the EPS mutant compared to wild-type cells before normalisation to cells (measured as chlorophyll) on the electrode. However, the amount of biofilm alone did not account for the greater photocurrent in the mutant after normalisation to chlorophyll loading (**Fig. 2b**). We therefore carried out further experiments to determine how the EPS affects photocurrent.

### Enhanced photocurrent output in the mutant with less exopolysaccharide strain is not due to salt stress

*Synechocystis* is a freshwater cyanobacterial species, and the EPS is known to confer protection to cells against stress caused by high concentrations of sodium chloride or the presence of toxic metal ions such as Cd^2+^ or Co^2+^, with the EPS-deficient strain used here growing more slowly than wild-type under those conditions.^28^ Previous studies have shown that cyanobacterial terminal oxidases affect the cells’ exoelectrogenic activity,^37^ and (for the cyanobacterium *Synechococcus elongatus* PCC7942) that the distribution of terminal oxidases between thylakoid and cytoplasmic membranes is influenced by the external concentration of sodium ions.^47^ It is also known that wild-type *Synechocystis* cells have increased ability to reduce exogenous ferricyanide under elevated salt concentrations.^38^ We therefore hypothesised that elevated sodium chloride concentrations might enhance cyanobacterial exoelectrogenic activity (which is not necessarily equivalent to ferricyanide reduction). If so, given that the EPS mutant cells lack the full protection afforded by EPS, these cells might be under mild salt stress even in BG11 medium, and this might account for some or all of the increased photocurrent from the EPS mutants.

To test if elevated sodium chloride concentrations lead to increased photocurrent output from *Synechocystis* we studied the effects of changes in concentrations in the electrolyte on photocurrent. The photocurrent profile and output of wild-type *Synechocystis* cells were first measured in standard BG11 electrolyte. Subsequently, additional NaCl or KCl were introduced into the BG11 electrolyte and the corresponding changes in photocurrent output after a 20-minute incubation period were measured. We also tested the effect of 600 mM sorbitol, to increase the osmotic potential without additional sodium or potassium ions. Addition of 600 mM NaCl to the BG11 electrolyte in fact resulted in a substantial decrease in photocurrent output from 280 ± 127 A cm^-2^ mol^-1^_Chl_ to 55 ± 50 A cm^-2^ mol^-1^_Chl_ (n = 3, P = 0.04) (**Supplementary Fig. 5**). Similarly, the addition of 600 mM KCl reduced the photocurrent output from 151 ± 18 A cm^-2^ mol^-1^ to 57 ± 39 A cm^-2^ mol^-1^ (n = 4, P = 0.002).

The addition of 600 mM sorbitol had no discernible effect on photocurrent output. The decrease in photocurrent from wild-type cells under salt stress indicates that the increased photocurrent generation by EPS mutant cells compared to wild-type was not due to an increase in sensitivity to salt in the EPS mutant cells.

### Evidence of a diffusible electron mediator in *Synechocystis sp*. PCC 6803

To test if diffusible species from IEET were important for the photocurrent output, convection was imposed by stirring the electrolyte to change the diffusion layer at the surface of the electrode. This approach has been widely used to assess the role of diffusible redox mediators.

Stirring the electrolyte at 1500 rpm diminished the photocurrents of wild-type cells significantly to 10 ± 4 % of that seen without stirring (n = 3, P = 0.004) (**Fig. 6**). [We also screened lower speeds, which reduced the photocurrents by similar amounts (**Supplementary Fig. 6**).] This provided further evidence for the involvement of a diffusible mediator in exoelectrogenesis in wild-type cells.

**Fig. 6.**
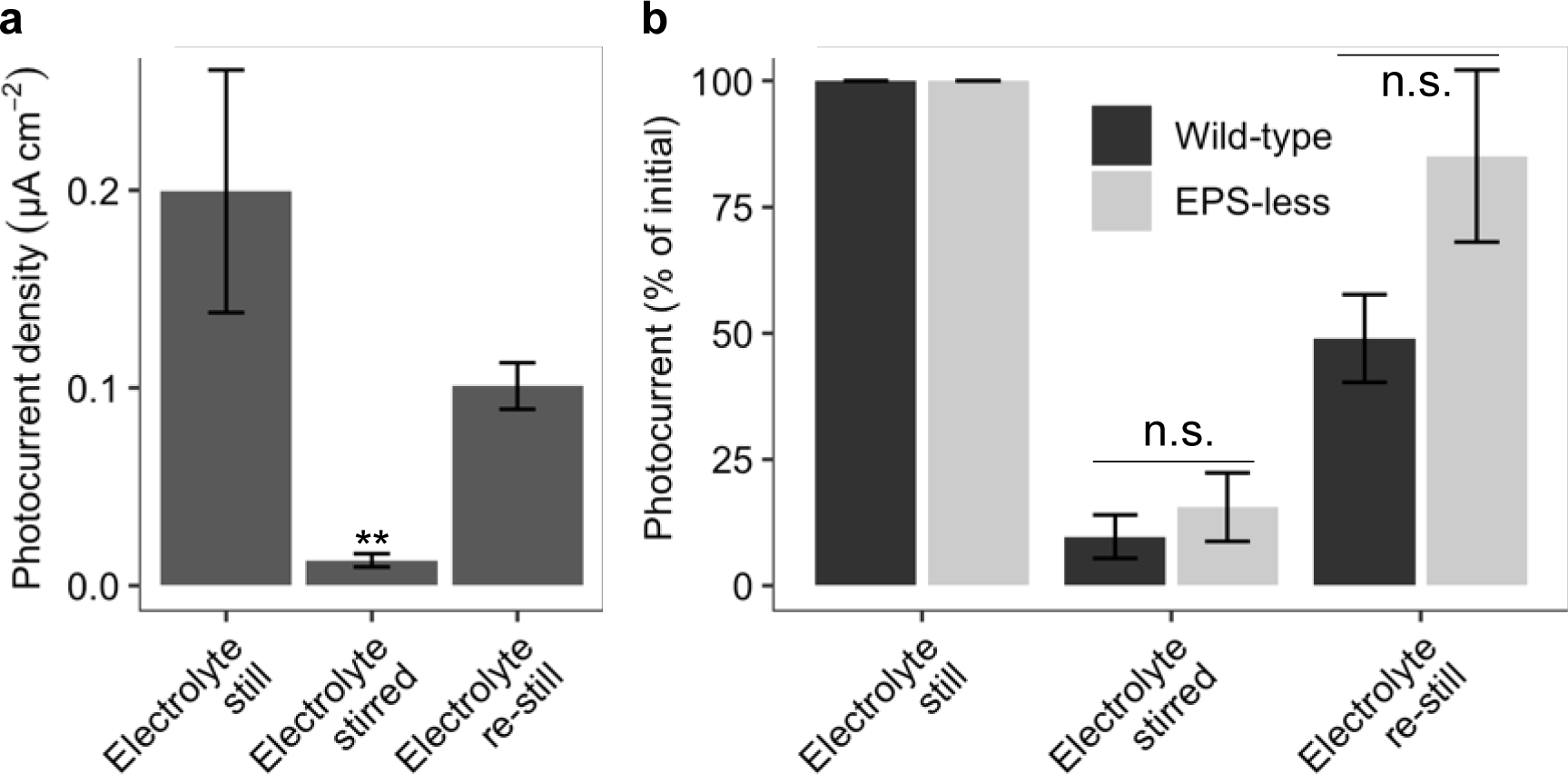
Effect of stirring the electrolyte. Photocurrent outputs with a magnetic stirrer bar stirring the BG11 electrolyte during chronoamperometry. Chronoamperometry was carried out under standardised conditions as described in Fig. 2. **a** Photocurrent outputs of wild-type cells. Data are presented as mean ± standard error of three biological replicates, ** = P < 0.01 compared to still electrolyte at the start of the experiment using a one-way ANOVA and Tukey HSD. **b** Photocurrent outputs of wild-type and EPS-less mutant cells. Data are presented as percentage of still electrolyte at the start of the experiment, mean ± standard error of three biological replicates, n.s. (not significant) = P > 0.05 compared between wild-type and EPS-less strains.

To test if redox mediators were also important for the EPS mutant, we repeated the experiment with it. Stirring the electrolyte diminished the photocurrents of EPS-less cells to 16 ± 7 % of that seen without stirring, which was not significantly different from the reduction in photocurrent from wild-type cells caused by stirring (**Fig. 6b**). Therefore, exoelectrogenesis in the EPS mutant also occurs predominantly through IEET. After stirring, the photocurrents of the EPS-less mutant cells returned closer to their starting levels (85 ± 17 %) than did the wild-type cells (49 ± 9 %), although the difference was again not statistically significant. The effects seen would be consistent with EPS-less mutant cells forming stronger biofilms than wild-type cells (**Fig. 3**), with less loss of cells under the experimental conditions.

### Effect of exopolysaccharide on consequences of adding small molecules

To test whether the presence of EPS affected the ability of molecules to diffuse in the microenvironment around the cells (and also to an electrode), we studied the interaction between cells and the dye carboxy-SNARF-acetoxymethylester (C-SNARF-AME).^48^ C-SNARF-AME equilibrates between a positively charged phenolic form and a neutral phenolate form, with a pKa of approximately 7.5 (**Supplementary Fig. 7**).^49^ It is expected to be predominantly in the phenolate form in the extracellular medium, with an increased proportion in the phenolic form in the cytoplasm when it has entered cyanobacterial cells.^50^ The forms have different fluorescence properties, allowing detection of dye that has entered the cells without interference from dye outside the cells.

A monolayer of cells was loaded into a microscopy dish and left to settle for 16 h, which was similar to the protocol for photoelectrochemistry experiments. Cells were treated with C-SNARF-AME and incubated for 30 min before imaging in the phenolic channel (λ_Ex_ = 540 nm, λ_Em_ = 550-590 nm), phenolate channel (λ_Ex_ = 590 nm, λ_Em_ = 610 – 630 nm), and the chlorophyll channel to determine cell boundaries (λ_Ex_ = 660 nm, λ_Em_ = 700 – 750 nm). Fluorescence values per unit area (as fluorescence intensity per pixel) were determined to take into consideration different cell sizes (**Fig. 5c**). For a negative control, cells were treated with BG11 medium with no fluorescent probe.

Without the addition of the C-SNARF-AME probe, there was very little signal in the phenolic channel inside either wild-type or EPS mutant cells (**Fig. 7**). This showed that chlorophyll autofluorescence should not interfere with the measurement of the phenolic form of C-SNARF-AME. (On the other hand, the phenolate form showed significant spectral overlap with chlorophyll autofluorescence (**Supplementary Fig. 8**).) Following incubation with the C-SNARF-AME probe, the intracellular phenolic fluorescence did not increase in wild-type cells (n = 3, P = 0.57), but increased approximately 6-fold in the EPS mutant cells (n = 3, P = 0.009) (**Fig. 7a,b**). No difference in the chlorophyll autofluorescence was observed between untreated EPS and wild-type cells (n = 3, P = 0.47), and the chlorophyll autofluorescence did not change after C-SNARF-AME treatment for either strain (**Supplementary Fig. 9**). These results suggest that EPS can inhibit diffusion and/or entry of some molecules to the cells.

**Fig. 7.**
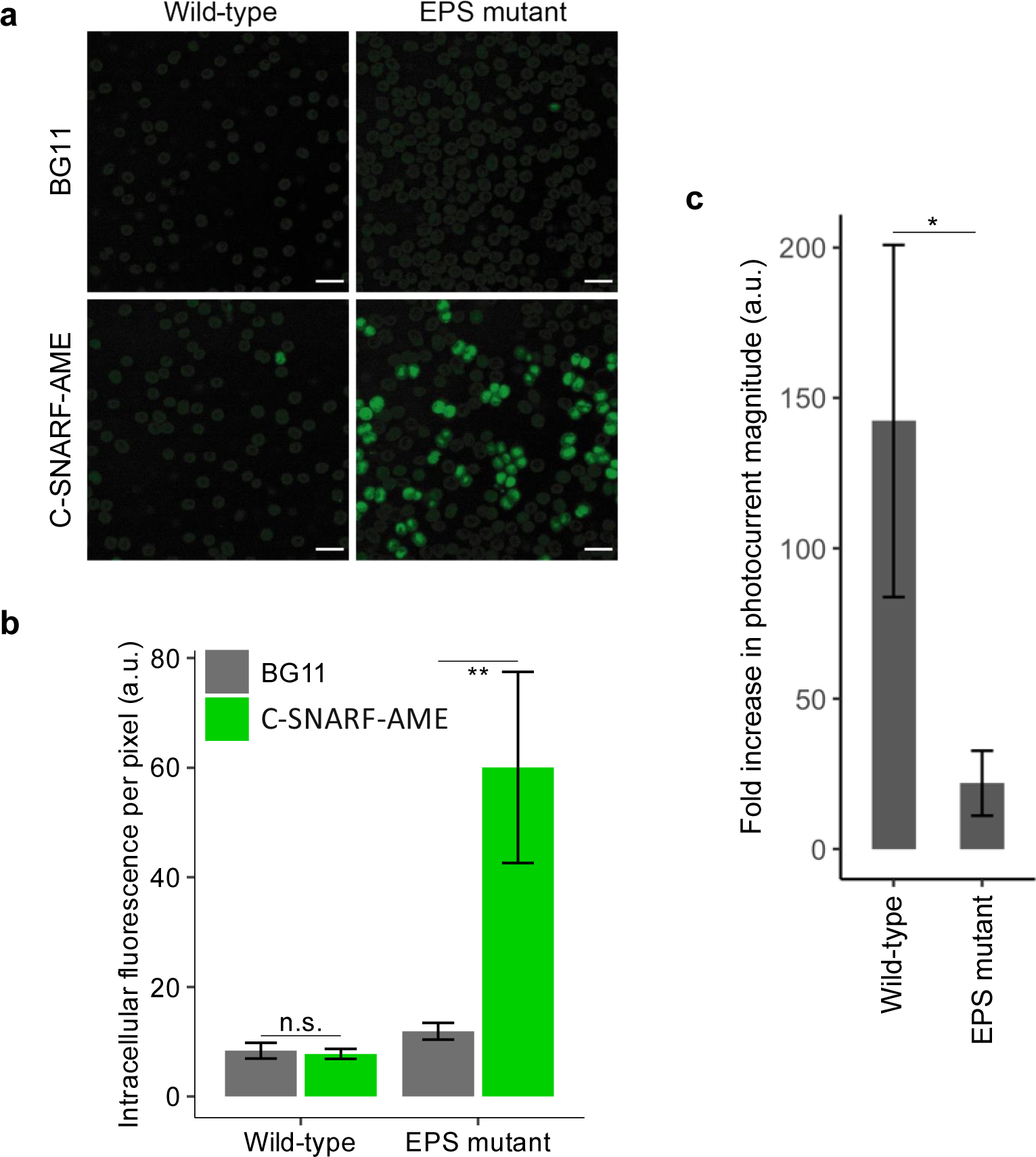
Consequences of exopolysaccharide for effects of small molecules. **a** Fluorescence microscopy images of wild-type and EPS mutant cells treated with BG11 medium or 10 µM C-SNARF-AME. Imaged in the phenolic channel: λ_Ex_ = 540 nm, λ_Em_ = 550-590 nm. Images representative of three biological replicates. Scale bar = 5 µm. **b** Intracellular phenolic C-SNARF-AME fluorescence normalised to cell area (by fluorescence intensity per pixel). Data presented as the mean ± standard deviation of 30 cells (3 biological replicates each measuring 10 cells, and errors propagated, n = 3 used for statistical analysis), n.s. (not significant) = P > 0.05, ** = P < 0.01 using a student’s t-test. **c** Photocurrent outputs from cells with DCBQ (1 mM) electron mediator are shown relative to their photocurrent output with no exogenous mediator added as 1 arbitrary unit (a.u.). Data are presented as mean ± standard deviation of three biological replicates, * = P < 0.05 compared to wild-type using a one-way ANOVA.

We reasoned that, if the EPS inhibits access of some molecules to the cell, the addition of an artificial redox mediator to enhance photocurrent might lead to a larger increase in photocurrent for the mutant strain than the wild-type. Chronoamperometry experiments were therefore performed in the presence of the electron mediator 2,6-dichloro-1,4-benzoquinone (DCBQ), with an applied potential of 0.5 V vs SHE for maximal mediation (DCBQ *E*_m_ = 0.315 V vs SHE).^9,24^ The increase in photocurrent output in the presence of DCBQ relative to the photocurrent without the exogenous mediator added was calculated. Surprisingly, although addition of DCBQ increased photocurrent for EPS mutant cells by approximately 20-fold, it gave rise to an approximately 140-fold increase for wild-type cells, which was significantly greater (n = 3, P = 0.011, **Fig. 7c**).

## Discussion

With similar amounts of cell material (assessed by the amount of chlorophyll) loaded on the electrodes initially, the EPS mutant cells showed four-fold greater photocurrent density than wild-type cells, although the kinetics of the current profiles were similar. *Synechocystis* mutants without EPS showed stronger biofilm formation on ITO-PET electrodes and more autoflocculation than wild-type cells. This may have helped cells pack better into the hierarchical IO-ITO electrodes, resulting in some of the increased photocurrent density. This is consistent with previous studies that show that disruption of EPS biosynthesis and/or export in other genes leads to flocculating strains with modulated adherence properties.^51,35,52^ For example, mutation of *slr0977* and *slr0982* genes that encode the permease and oligosaccharide binding proteins that function together as a ABC-transporter resulted in increased biofilm formation on glass in monolayers.^51^ The enhanced biofilm formation of the EPS deficient mutant may have been a consequence of its less negative cell surface charge (negative zeta potential) and cell surface hydrophobicity.^28^

These results support previous suggestions that attempts to increase BPV outputs should optimise the cell-electrode interface, including increasing cell adherence to the electrode. Previously a hyperpiliated Δ*pilT1* mutant was shown to have significantly enhanced biofilm formation on ITO-PET than mutants with fewer thick pili, indicating that the type IV pili are involved in cell-electrode adherence.^9^ Manipulation of the EPS is also an attractive possibility for modulating the cell surface charge and electrode adherence, and it will be interesting to explore the effects of combining hyperpiliation and deprivation of EPS.

After normalising for the amount of chlorophyll in material retained on the electrode, the EPS mutant still showed double the photocurrent density. Therefore, the EPS, either itself or as a scaffold for other redox active molecules, is not involved in a DEET mechanism of exoelectrogenesis in *Synechocystis*. This is also the case for type IV pili in *Synechocystis*.^8,9^

Elevated NaCl concentrations were shown to decrease current output from wild-type cells, indicating that the increased current seen in the EPS mutant was not due to an increased sensitivity to NaCl in their cellular microenvironment in the standard BG11 medium. The EPS mutant cells have been shown to grow at a similar rate to wild-type cells in standard BG11 medium,^28^ showing the viability of this mutant for use in biotechnologies under appropriate conditions. By contrast, many of the other cyanobacterial strains shown to have enhanced current output also show impaired growth or viability.^10,37,53,54^

Experiments using stirred systems indicated that much of the photocurrent detected was dependent on a diffusible electron mediator, consistent with existing indications of an endogenous electron mediator in cyanobacteria.^24–26,10^ Fluorescence microscopy experiments with a fluorescent probe demonstrated that the EPS can hinder the ability of small molecules to enter the *Synechocystis* cell. If the EPS hinders the movement of the diffusible mediator, this might contribute to the increased exoelectrogenic activity of the EPS mutant.

Addition of the exogenous electron mediator DCBQ led to a much greater increase in electron export from wild-type cells than from the EPS mutant. The interpretation of this is complex, as a number of effects might be proposed. The enhancement caused by DCBQ might be due to its ability to extract electrons from photosystems soon after photo-excitation,^55^ in which case the smaller enhancement seen with EPS mutant cells could be due to off-target inhibitory effects caused by increased entry of DCBQ in the absence of EPS. (A decrease in current mediation by DCBQ at concentrations around or above 1 mM has been reported,^55^ though the reason is not clear.) DCBQ might also enhance electron output from some other component(s) in the photosynthetic electron transfer chain or the cytosol, able to drive electron export in the absence of DCBQ, but with export enhanced by DCBQ. In that case, the smaller increase seen with the EPS mutant might again be explained by off-target effects of DCBQ in the mutant. Alternatively, if the existing sites from which DCBQ-facilitated electron transfer were more efficiently coupled (‘wired’) to the extracellular environment in the EPS cells than in the wild-type in the absence of DCBQ, a more modest effect of DCBQ addition would be expected in the EPS mutant cells. Further analysis of the action of DCBQ will be helpful.

## Conclusion

We conclude that indirect extracellular electron transfer is the bioelectrochemical mechanism of exoelectrogenesis in *Synechocystis*. We propose a model (**Fig. 8**) for the EPS-deficient cells whereby significant reduction of EPS leads to the formation of biofilms on electrodes with greater cell density, stronger electrode adherence and increased photocurrent (even after normalisation to the amount of chlorophyll present). The reduction of EPS means that small molecules, which may include endogenous electron mediators, are more easily able to access the cells. Overall, exopolysaccharide deprivation enhances photocurrent generation of *Synechocystis*. However, added exogenous mediators may have a less beneficial effect on overall current production by EPS cells than on current production by wild-type cells. Use of EPS-deficient strains is likely to offer further enhancement of power output from photosynthetic microorganisms in practical applications of BPVs in the long term, especially where the use of expensive artificial mediators with potentially cytotoxic side-effects is undesirable.

**Fig. 8.**
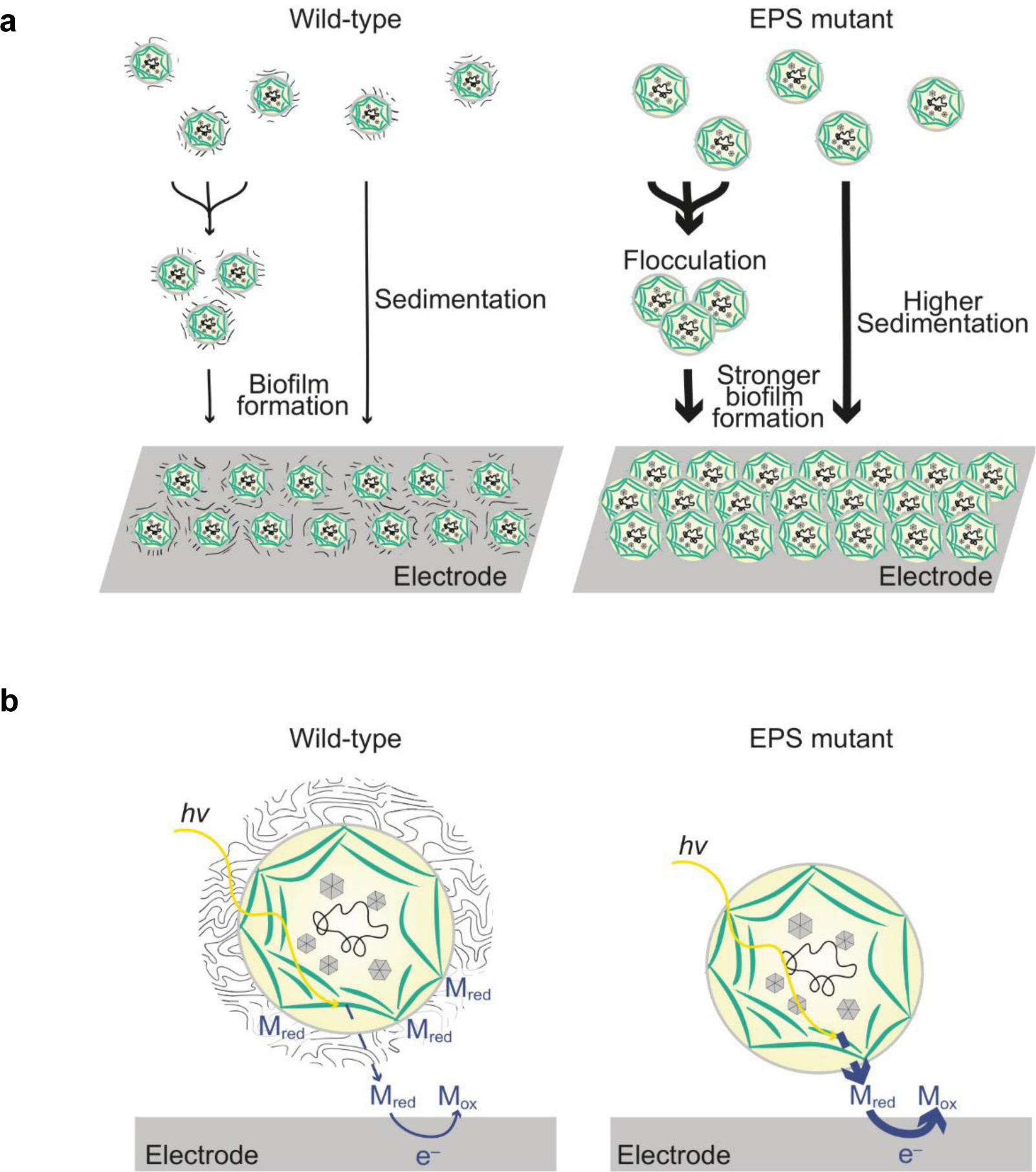
Proposed model of the role of exopolysaccharides in exoelectrogenesis. **a** Removal of the exopolysaccharides (EPS) increases flocculation, sedimentation, and biofilm formation, leading to increased biocatalyst loading on electrodes. **b** Removal of the EPS decreases the distance that endogenous electron mediators need to diffuse to reach the electrode and allow more rapid diffusion, leading to improved electronic wiring of cells to the electrode.

## Materials and Methods

### Cell Culture and Strains

The *Synechocystis sp.* PCC 6803 EPS mutant and background wild-type strain were a gift from Prof. Franck Chauvat (Institut de Biologie Intégrative de la Cellule (I2BC), France). The EPS mutant has the *slr1875* and *ssll1581* genes interrupted with a kanamycin resistance cassette and a spectinomycin resistance cassette respectively.^28^

Cyanobacterial cells were cultured photo-autotrophically under 50 µmol_photons_ m^-2^ s^-1^ of continuous white light at 30°C in BG11 medium supplemented with 10 mM NaHCO_3_.^56^ Liquid cultures were bubbled with air and shaken at 120 rpm. 1.5% (w/v) agar was used in solid medium. Antibiotics were added at a final concentration of 50 µg ml^-1^ kanamycin and 2.5 µg ml^-1^ spectinomycin for routine culture of the EPS mutant, but not for preparing EPS mutant cells for experiments where they were compared with the background wild-type strain. Culture growth was measured by attenuance at 750 nm (OD_750_). Chlorophyll concentrations of cultures (nmol_Chl_ ml^-1^) were calculated from absorbances at 680 nm and 750 nm: (A_680_-A_750_)×10.814.^53^ All measurements were taken using a UV-1800 Spectrophotometer (Shimadzu).

### Photoelectrochemistry

All photoelectrochemical measurements were performed using an Ivium Technologies CompactStat, in BG11 medium (pH 8.5) as the electrolyte, at 25°C, under atmospheric conditions, with an Ag/AgCl (saturated KCl) reference electrode (corrected by + 0.197 V for SHE), a platinum mesh counter electrode and cyanobacterial cells loaded on inverse opal ITO (IO-ITO) working electrode. IO-ITO electrodes with 10 μm macropores and 3 μm interconnecting channels at a thickness of 40 µm were prepared using templated synthesis via an infiltration approach as previously described.^18^ Planktonic cultures of early stationary phase cyanobacterial cells at OD_750_ of ca. 1 were loaded onto the IO-ITO electrodes as previously described.^9^

Chronoamperometry experiments were performed at an applied potential 0.3 V vs SHE, or 0.5 V vs SHE in the presence of 1 mM DCBQ exogenous mediator that had been added to the electrolyte for 15 min in darkness beforehand. Chronoamperometry experiments were performed at a sampling rate 1 s^-1^, with 60s/90s light/dark cycles using a collimated LED (50 µmol_photons_ m^-2^ s^-1^ λ_max_ = 680 nm, approximately 1 mW cm^-2^ equivalent).

For photoelectrochemistry experiments where the electrolyte was stirred, the static three-electrode photoelectrochemical set up was placed a few mm above a stirrer plate. A 4 mm long, rod-shaped, smooth magnetic stirrer bar was placed in the electrolyte. The stirrer plate was switched on during photoelectrochemistry experiments to stir the electrolyte.

The current outputs were normalised to the geometric area of the IO-ITO electrode (0.79 cm^2^) to obtain current densities. The photocurrent output was calculated as the difference between the steady state currents in the light (mean current output in the final 5 seconds of the light period) and dark (mean current output in the final 5 seconds of the dark period). The chlorophyll content of the loaded IO-ITO electrode was determined after photoelectrochemistry experiments using a methanol extraction as previously reported,^9^ and photocurrent outputs were normalised by chlorophyll loading. The photocurrent enhancement with DCBQ added was calculated relative to the photocurrent without an exogenous mediator added.

### Scanning electron microscopy

Cyanobacterial cells were loaded onto IO-ITO electrodes and left for a set amount of time under darkness as per the protocol for photoelectrochemistry. The loaded electrodes were rinsed in BG11. The loaded electrodes were then fixed in liquid nitrogen-cooled ethane and freeze-dried overnight. The loaded electrodes were mounted on aluminium stubs with silver dark and coated with 15 nm iridium using an EMITECH K575X Peltier cooler. The sample was stored in a desiccator until imaged using a TESCAN MIRA3 FEG-SEM with a 30 kV beam acceleration. Images were processed using Illustrator.

### Biofilm formation assay

A crystal violet assay was used to measure biofilm formation as previously reported.^9^ Cyanobacterial cells were loaded onto ITO-coated polyethylene terephthalate (ITO-PET) 1 cm^2^ discs in wells of a 24-well plate and left under darkness as per the protocol for photoelectrochemistry. 150 µl was aspirated from the anodic chambers without disturbing the biofilm before gently transferring the cell-loaded discs to the wells of a 24-well plate. The cell-loaded discs were gently washed thrice with 750 µl BG11 medium to remove non-adherent cells. Crystal violet solution (250 µl, 0.1% w/v) in dH_2_O was pipetted into each well and incubated for 15 min at room temperature. Following this, 200 µL was aspirated from each well and the disc was washed three times with 250 µl dH_2_O. 1 ml DMSO was pipetted into each well and the 24-well plate was vigorously shaken for 15 min. The absorbance at 600 nm of the crystal violet in the DMSO supernatant was measured and used as a proxy for measuring the amount of cellular material bound to the ITO-PET.^57^

### Flocculation assay

Cell-cell interaction was measuring using a flocculation assay adapted from a protocol reported previously.^46^ Exponential phase cultures were diluted to an OD750 of 0.5 and early stationary phase cultures were diluted to an OD750 of 1.0. Wells of a 6-well plate were filled with 5 ml of culture. The wells were sealed and incubated for 48 h at 30°C under 40 µmol_photons_ m^-2^ s^-1^ light with shaking at 75 rpm. The plates were inspected visually and photographed post incubation. Images were processed for a quantitative measurement of flocculation (Supplementary Fig. 3). The images were converted to black and white and cropped to include only the central part of the well to remove light reflections on the periphery. Images were processed in R using the package magick. At each pixel the difference between the background and local light intensity was taken. The sum of all pixels where intensity was less than background was taken and divided by the number of pixels to control slight resolution variations across photographs of wells. The final value quantifying flocculation was the mean of all intensities in the image darker than the image background, where the image background is the mean of intensities of all pixels in the entire image.

### Fluorescence microscopy and dyes

Equal volumes of cell culture (200 µl) at early stationary phase (OD_750_ = 1.2) were plated onto an ITO-coated glass coverslip fixed to a microscopy dish. Cells were left for 16 hours in the dark to settle and form a biofilm. Brightfield images were taken using an EVOS M5000 epifluorescence microscope, using a 63X objective. Cell diameter was calculated by selecting 10 random cells each in three biological replicates and counting the number of pixels across either vertically or horizontally. The number of pixels across was then multiplied by 107 nm per pixel. The same approach was used to determine the distance between cells.

For the probe experiments, cells were similarly left for 16 hours in the dark to settle and form a biofilm. Fluorescence images were taken using a Leica Stellaris 5 Confocal Microscope, using a 63X objective at either 1X (for images used for quantification) or 4X (for representative images) digital zoom. Thirty minutes prior to imaging, 5 mL of either BG11 (negative control) or 10 µM C-SNARF-AME in BG11 were added to the microscopy dish. Images were then taken in the following channels: Chl *a*, l_Ex_ = 660 nm, l_Em_ = 700 – 750 nm; C-SNARF-AME (phenolic), l_Ex_ = 540 nm, l_Em_ = 550-590 nm; C-SNARF-AME (phenolate), l_Ex_ = 590 nm, l_Em_ = 610 – 630 nm. Intracellular fluorescence quantification was calculated by selecting 10 random cells each in three biological replicates, tracing their outline and quantifying their average pixel intensity using the Measure function in ImageJ.

For cell dimension quantifications (cell-cell distance, cell diameter, cell diameter for normalising fluorescence), the mean and standard deviation were calculated for 10 cells in each biological replicate, then mean of means taken, and error propagation calculated across the three groups for the standard deviation. A sample size of n = 3 was used for statistical analysis.

## Acknowledgements

This work was supported by the Cambridge Trust (to L.T.W. and L.S.), UK Research and Innovation (BB/R011923/1 to J.Z.Z., BB/M011194/1 to E.W.), China Scholarship Council (to L.S.), and the Novo Nordisk Foundation (NNF22OCOO79717 to L.T.W.). The *Synechocystis sp.* PCC 6803 EPS mutant and background wild-type strain were a gift from Prof. Franck Chauvat (Institut de Biologie Intégrative de la Cellule (I2BC), France). The differently sized wild-type strains Arizona and Imperial were a gift from Prof. Conrad Mullineaux (Queen Mary University of London, UK).

## Author contributions

**L.T.W.:** Conceptualisation, Investigation, Visualization, Writing - Original Draft, Writing - Review & Editing, Supervision, Funding acquisition. **E.W.:** Methodology, Investigation, Writing - Review & Editing. **V.S.:** Investigation, Software, Visualization, Writing - Original Draft. **L.S.:** Investigation. **X.C.** Investigation. **J.Z.Z.** Methodology, Resources, Writing - Review & Editing, Supervision, Funding acquisition. **C.J.H.:** Resources, Writing - Original Draft, Writing - Review & Editing, Supervision, Funding acquisition, Project administration.

