## Supplementary Information for "Fourfold increase in photocurrent generation of *Synechocystis* sp. PCC 6803 by exopolysaccharide deprivation"

### *Synechocystis* sp. PCC 6803 by exopolysaccharide deprivation

### Supplementary information

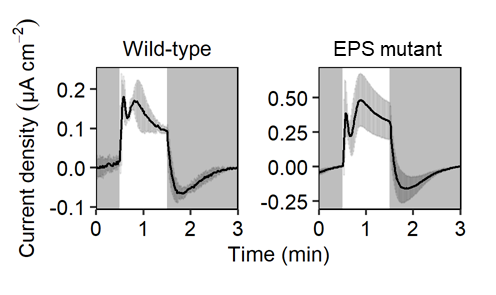

Supplementary Fig. 1 Photocurrent profiles.

Data are the same as in Fig. 2A, but with different y axes. White panels = light on and grey panels = light off. The current was normalised to the geometric area of the IO-ITO electrode to obtain current densities. Data are presented as mean ± standard deviation of three biological replicates.

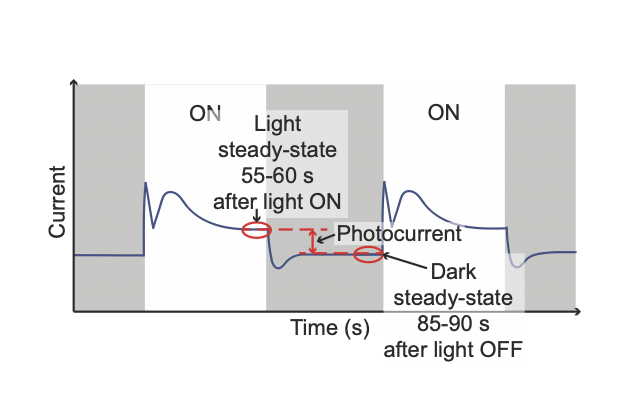

Supplementary Fig. 2 Analysis of photocurrent profiles.

Cartoon of a chronoamperometric trace of cyanobacterial cells on an IO-ITO electrode showing light/dark cycles annotated to show the steady state current in the light, steady state current in the dark and the photocurrent.

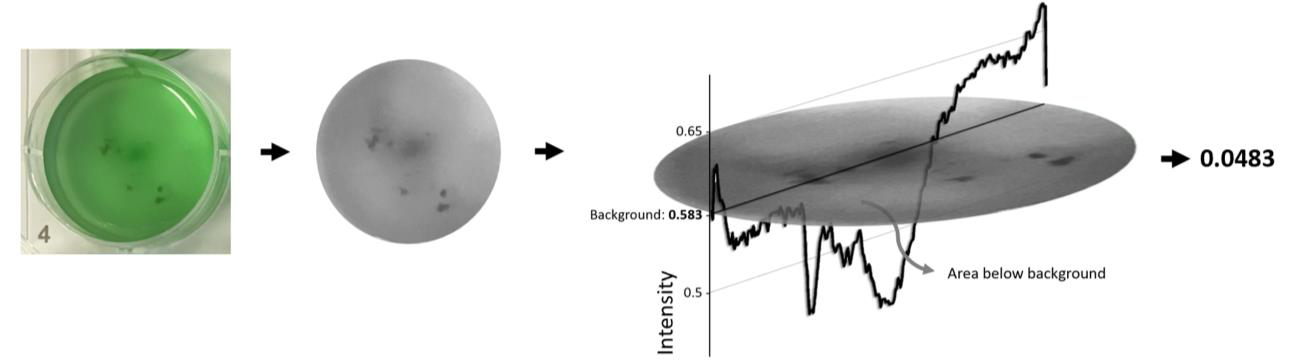

Supplementary Fig. 3 Image processing algorithm.

The images were converted to black and white and cropped. At each pixel the difference between the background and local light intensity was taken.

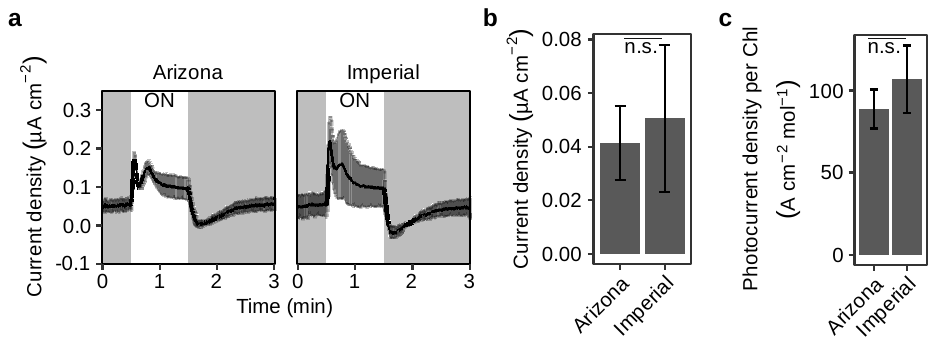

Supplementary Fig. 4 Photoelectrochemical performance of two *Synechocystis* wild-type strains does not depend on cell size. The figure shows a Photocurrent profiles, b photocurrent outputs, and c photocurrent outputs normalised to chlorophyll on electrode after photoelectrochemical measurements. Chronoamperometry under standardised conditions as described in Fig. 2. Results presented as the mean of biological replicates (Arizona: n = 2, Imperial: n = 3), ribbons/error bars are standard error of the mean, P > 0.05 from one-way ANOVA indicates not significantly different (n.s.). The ‘Imperial’ strain has approx. 1.4x greater diameter than the ‘Arizona’ strain.(LTW unpublished and Ref. ^1^)

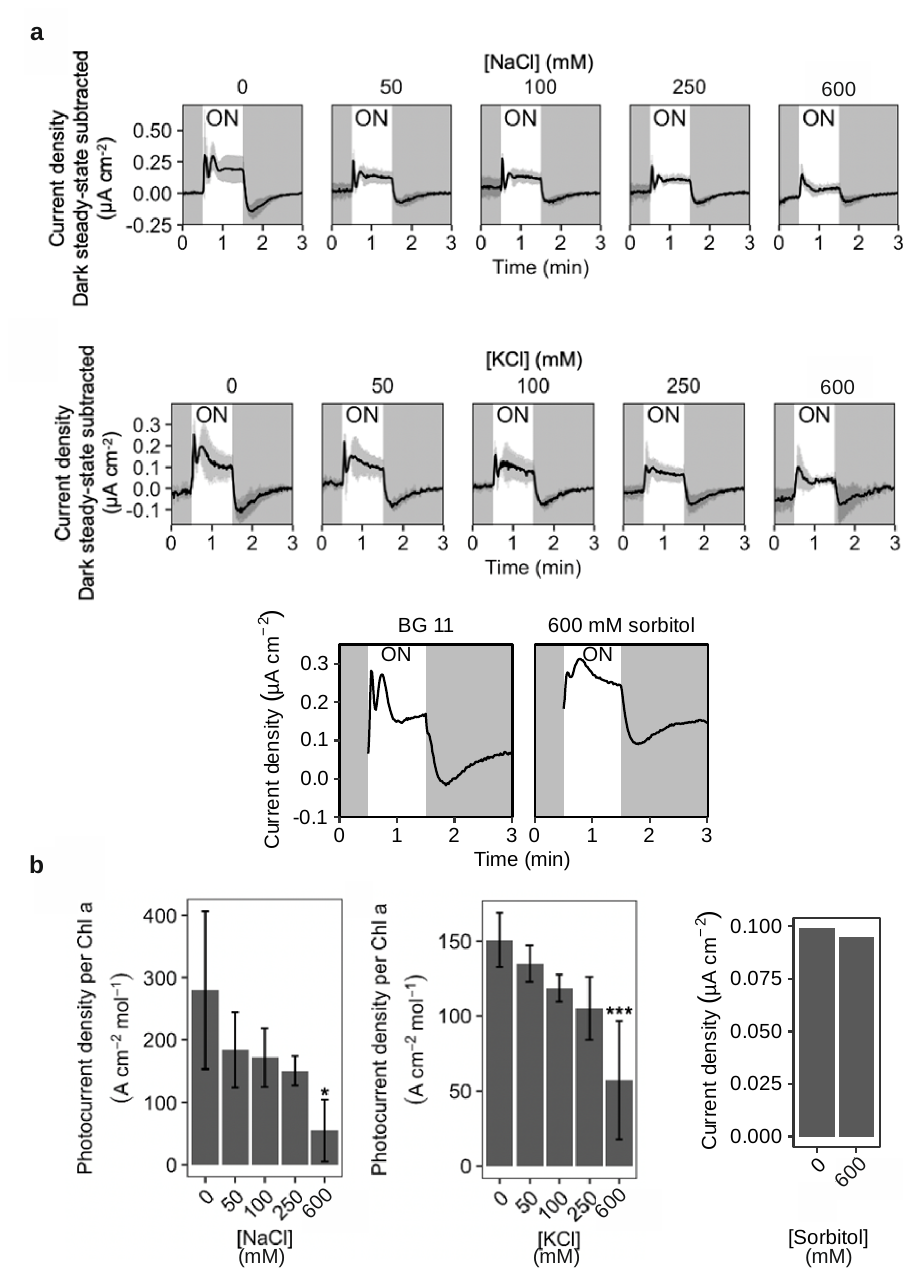

Supplementary Fig. 5 Effect of adding extra salt or osmolyte to the BG11 electrolyte.

Wild-type *Synechocystis* cells in standard BG11, and BG11 with increasing concentrations of NaCl, KCl or sobitol in the electrolyte. **a** Photocurrent profiles, **b** Photocurrent outputs (normalised for chlorophyll loading if replicates). Chronoamperometry under standardised conditions as described in Fig. 2, apart from ion concentrations specified. Data presented as the mean (solid black line in photocurrent profiles) ± standard deviation (grey bars in photocurrent profiles) of three biological replicates (sorbitol: n =1), * = P < 0.05 by one-way ANOVA compared to standard BG11. ON = light on.

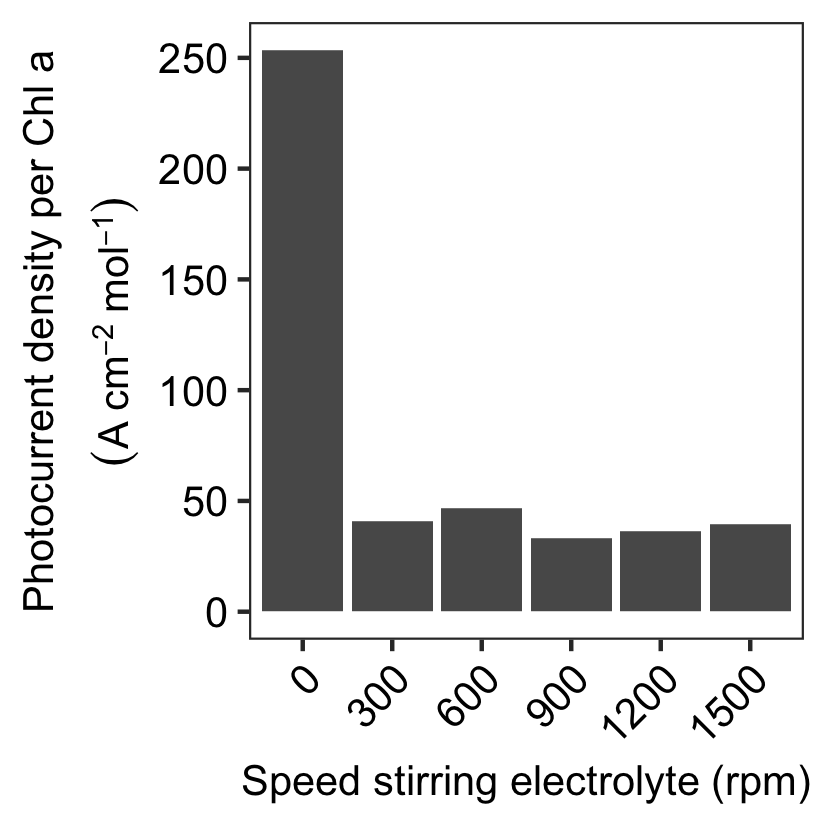

Supplementary Fig. 6 Effect of stirring the electrolyte at different speeds.

Photocurrent outputs of wild-type *Synechocystis* cells in BG11 electrolyte stationary or with stirring at different speeds, normalised to chlorophyll loading. Chronoamperometry under standardised conditions as described in Fig. 2. One biological sample is shown.

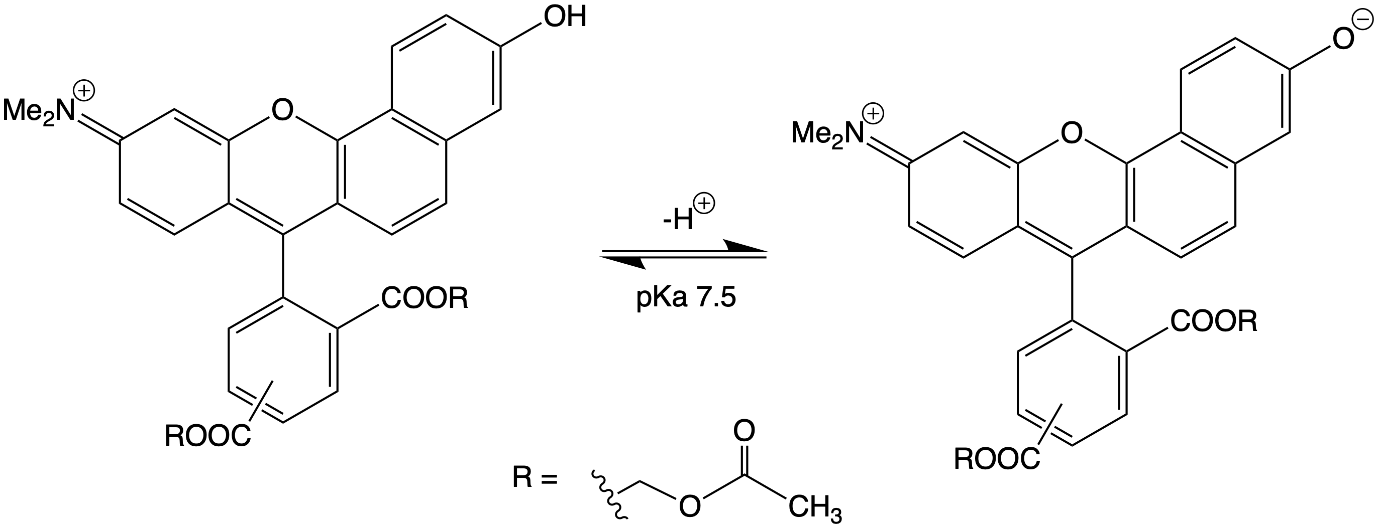

Supplementary Fig. 7 Schematic showing C-SNARF-AME structure and deprotonation. The pH probe carboxy-SNARF-acetoxymethylester exists in equilibrium between its phenolic (left) and phenolate (right) forms, with a pKa of 7.5.

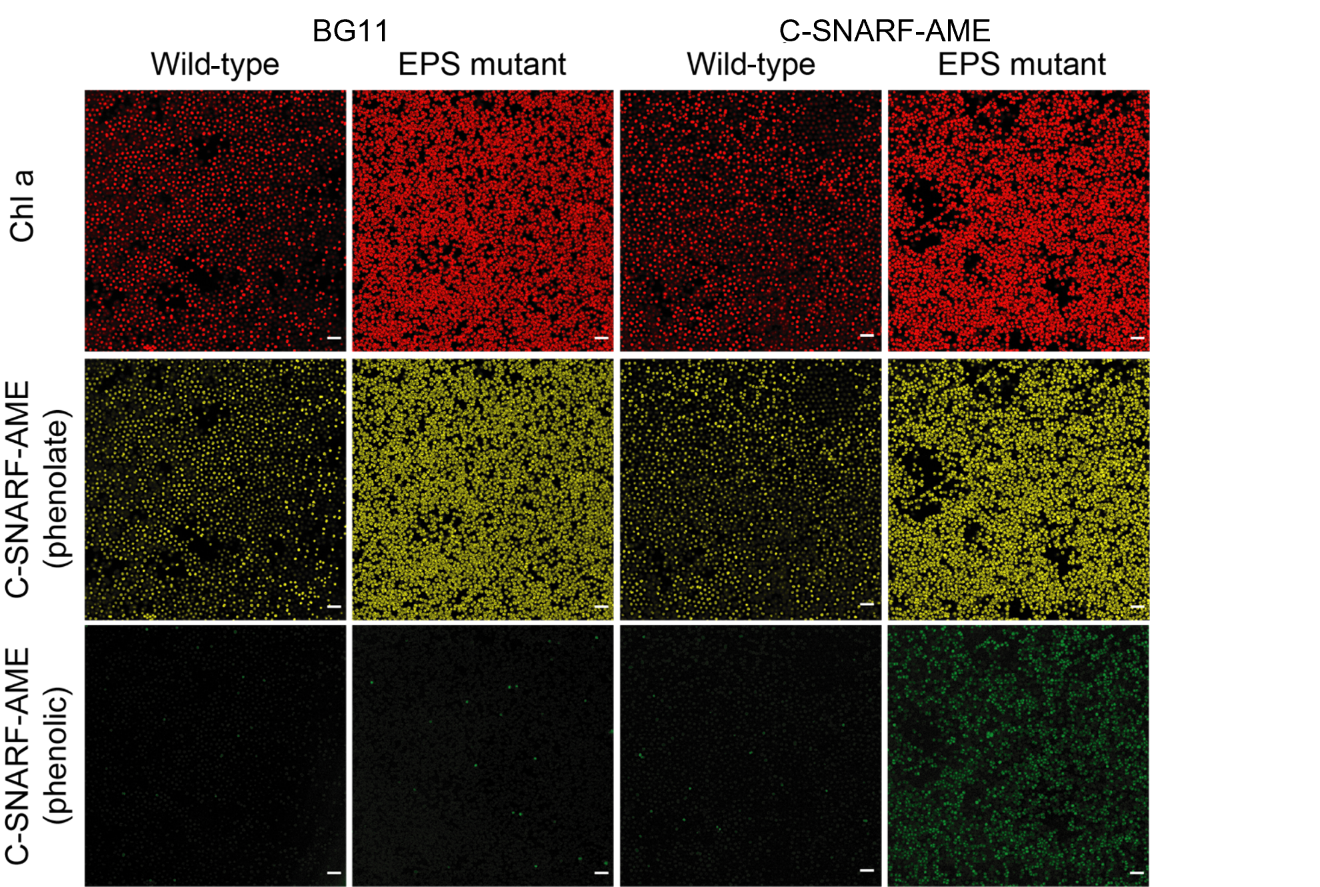

Supplementary Fig. 8 Fluorescence microscopy of wild-type and exopolysaccharide mutant cells with charged fluorescent probes.

Fluorescence microscopy images of wild-type and EPS mutant cells treated with BG11 medium or 10 µM C-SNARF-AME. Imaged in the Chlorophyll *a* (Chl a) channel: λ_Ex_ = 660 nm, λ_Em_ = 700 – 750 nm; phenolate channel: λ_Ex_ = 590 nm, λ_Em_ = 610 – 630 nm; and the phenolic channel: λ_Ex_ = 540 nm, λ_Em_ = 550-590 nm. Images representative of three biological replicates. Scale bar = 10 µm.

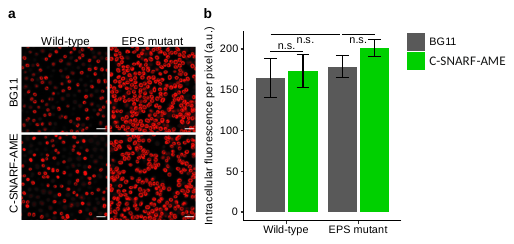

Supplementary Fig. 9 Chlorophyll autofluorescence microscopy images of biofilms of cells.

**A)** Fluorescence microscopy images of wild-type and EPS mutant cells treated with BG11 medium or 10 µM C-SNARF-AME. Imaged in the chlorophyll *a* channel: λ_Ex_ = 660 nm, λ_Em_ = 700 – 750 nm. Images representative of three biological replicates. Scale bar = 5 µm. **B)** Intracellular chlorophyll *a* fluorescence normalised to cell area (by fluorescence intensity per pixel). Data presented as the mean ± standard deviation of 30 cells (3 biological replicates each measuring 10 cells, and errors propagated, n = 3 used for statistical analysis), n.s. (not significant) = P > 0.05 using a student’s t-test.
